# Pre-adaptation to climate change through topography-driven evolution of traits and their plasticity

**DOI:** 10.1101/821561

**Authors:** H. De Kort, B. Panis, S.B. Janssens, K. Helsen, O. Honnay

## Abstract

Climate change is expected to increase the level of drought stress experienced by many plant populations, yet the spatial distribution of changes in dryness remains highly uncertain. Species can, to some extent, adapt to climate uncertainty through evolving increased trait plasticity. Biodiversity conservation could capitalize on such natural variation in the ability of populations to cope with climate variability. Yet, disentangling evolution of trait means vs. trait plasticity is challenging, as it requires a sampling design with genetic replicates grown under distinct environmental conditions. Here, we applied different soil moisture treatments to clones of Fragaria vesca plants that were raised from seeds that were sampled in distinct mountainous topographical settings, to study adaptive trait and plasticity divergence in response to drought. We demonstrate that various fitness traits evolved along topographical gradients, including increased specific leaf area (SLA) with increasing slope, and increased growth plasticity with increasing altitude. Our results indicate that traits and their plasticity can evolve independently in response to distinct topographical stressors. We further show that trait heritability varies considerably among traits and topographical settings. Heritability of phenotypic plasticity tended to increase with altitude for all traits, with populations from high altitudes harboring more than twice the heritability for growth and SLA plasticity compared to populations from low altitudes. We conclude that (i) low altitudinal populations, which are expected to be least vulnerable to climate change, may only withstand limited increases in drought stress, while (ii) populations that evolved to thrive under more heterogeneous mountain conditions are pre-adapted to climate change through high plasticity and heritability. Highly heterogeneous landscapes may thus represent invaluable sources of quantitative genetic variation that could support conservation under climate change across the globe.

## 1 INTRODUCTION

Observed and projected changes in temperature and precipitation are destabilizing existing interactions between plant and animal communities and their abiotic context across the globe (Urban *et al.* 2015; Bertrand *et al.* 2016; Vázquez *et al.* 2017; Alexander *et al.* 2018). While drought stress levels will continue to change worldwide with high confidence, the regional and local spatial distribution and magnitude of these changes remains highly uncertain (IPCC 2014). This uncertainty complicates biodiversity conservation anticipating climate change, for example through assisted migration of genotypes pre-adapted to projected climate conditions. Yet, conservation strategies aiming to mitigate climate change impacts in the absence of accurate projections could capitalize on natural variation in the ability of populations to cope with climate variability and uncertainty. Populations that have evolved in highly heterogeneous and temporarily variable environments likely retain a higher fitness for a wide range of environmental conditions than populations that are adapted to more homogeneous and stable environments (Reed *et al.* 2010; Chevin & Lande 2011; Chevin & Hoffmann 2017; Bonamour *et al.* 2019).

Established variation in phenotypic traits that underlie drought tolerance can greatly increase the ability of populations to withstand unpredictable changes in soil moisture levels. While plastic phenotypic variation underlying drought tolerance may support fitness in environments currently featured by heterogeneous soil humidity levels, heritable phenotypic variation is required for contemporary evolution towards an increase in drought tolerance and/or phenotypic plasticity for drought tolerance (Anderson *et al.* 2011; Palacio-López *et al.* 2015; Hoffmann *et al.* 2017; Kingsolver & Buckley 2017). Genotypes that are capable of adjusting their phenotypes to changing soil moisture levels through plastic responses in particular represent a valuable yet understudied component of the evolutionary potential of natural populations in the context of climate change anticipated drought events (IPCC 2014; van Kleunen & Fischer 2005; Kingsolver & Buckley 2017; Bonamour *et al.* 2019).

How particular environmental stressors drive evolution toward increased phenotypic plasticity in a population (i.e. increase the frequency of genotypes with enhanced levels of phenotypic plasticity) is a particularly challenging question, as answering it requires genetically identical replicates to be raised under distinct environmental settings (De Kort *et al.* 2016; Arnold *et al.* 2019). Although quite some research relied on genetically similar individuals to study plasticity evolution approximately (e.g. Sultan 2001; Zhang *et al.* 2012; Scheepens *et al.* 2018), to our knowledge, only one study has hitherto assessed how genetic variation in phenotypic plasticity varies across plant populations adapted to distinct climatic environments using genetically identical clones (Cooper *et al.* 2019). Transplantations of genotypic replicates from 16 populations of *Populus fremontii* to cool, warm and hot sites in the Southwestern United States revealed that genotypes originating from low latitude populations were significantly more plastic for bud set and bud flush as compared to genotypes originating from higher latitudes (Cooper et al. 2019). While this study provides rare evidence for natural variation in the magnitude of plastic responses to climate, the potential of natural populations to evolve this plasticity remains elusive. Elucidating the natural occurrence and heritability of trait plasticity evolution could thus shed renewed light on the ability of populations to cope with environmental changes. Replication within genotypes across treatments therefore represents a powerful experimental approach for (i) disentangling plastic (i.e. within-genotype) trait variation from genetic (i.e. among-genotype) trait variation, and (ii) studying heritable variation underlying trait plasticity.

Fine-scale environmental variation has frequently been associated with high levels of adaptive phenotypic variation. In particular, the highly heterogeneous landscapes from mountain ranges provide ample opportunity for adaptive radiation at fine spatial scales (Halbritter *et al.* 2018; Waterhouse *et al.* 2018). Moreover, high-altitudinal secondary valleys and cold air sinks typical of topographically complex landscapes designate topographical factors other than altitude as contributing determinants of temperature and soil moisture levels (Körner 2007; Günther *et al.* 2016; O’Brien *et al.* 2017; Pfennigwerth *et al.* 2017). For natural populations, fine-scale phenotypic variation coinciding with topographical variation could thus be key to cope with sudden environmental changes.

In the face of uncertain climate and soil moisture changes, the topographical complexity of mountainous landscapes may harbor an invaluable source of variation in drought resistance traits, including increased water use efficiency (e.g. through more and smaller stomata, Dittberner et al. 2018), lower specific leaf area (SLA, Bansal et al. 2015) and advanced flowering (Wilczek *et al.* 2010). In this regard, south-oriented slopes are assumed to be dominated by drought-resistant phenotypes, especially where strong inclinations facilitate water efflux. Populations inhabiting such challenging environments may thus evolve an enhanced ability to cope with drought stress through plastic or genetic phenotypic responses.

Here, we study topography-driven adaptive variation underlying drought stress-related traits and their plasticity, using genetic replicates of natural *Fragaria vesca* populations. A total of ca. 12 genotypes × 11 mountainous populations × 3 soil moisture treatments × 4 clones was monitored in a common garden for growth, vegetative propagation, SLA, stomatal density and flowering rate. Traits and their plasticity (quantified as relative distance plasticity indices (RDPIs)) were subsequently modeled in relation to slope, aspect, topographical position index, absolute altitude, and relative altitude of the population sampling locations. Heritability was calculated for each trait and for each RDPI. This methodological strategy allowed us to tackle the following research questions: (i) how much adaptive variation do mountainous populations harbor for traits (and their plasticity) related to drought resistance? (ii) are topographical variables other than altitude strong determinants of adaptive trait variation, and (iii) does trait plasticity, measured as RDPI, have a strong heritable component? In parallel, we hypothesized that genotypes originating from low absolute altitude (~ high temperatures), high relative altitude and high topographic position index (~ high water efflux) and steep, southerly oriented slopes (~ high water efflux and evapotranspiration) have evolved traits that allow to cope with drought stress.

## 2 METHODS

### 2.1 Study species and sampling

*Fragaria vesca* L., also known as the woodland strawberry, is a self-compatible, insect-pollinated perennial stoloniferous herb with a circumglobal temperate distribution. It is typically found along mixed and broadleaf forest edges, gaps and paths. Seeds from a total of eleven natural populations were sampled in the French Alps and Pyrenees, covering various topographical conditions. The populations were spatially clustered in four meta-populations (Fig. 1, Table 1), and distances between populations within a meta-population ranged from 550 m to 3700 m. This spatial clustering can be assumed to result in substantially stronger genetic relatedness within than between meta-populations. To minimize the risk of sampling the same genet twice, distance among samples was maximized within locations.

**Fig. 1.**
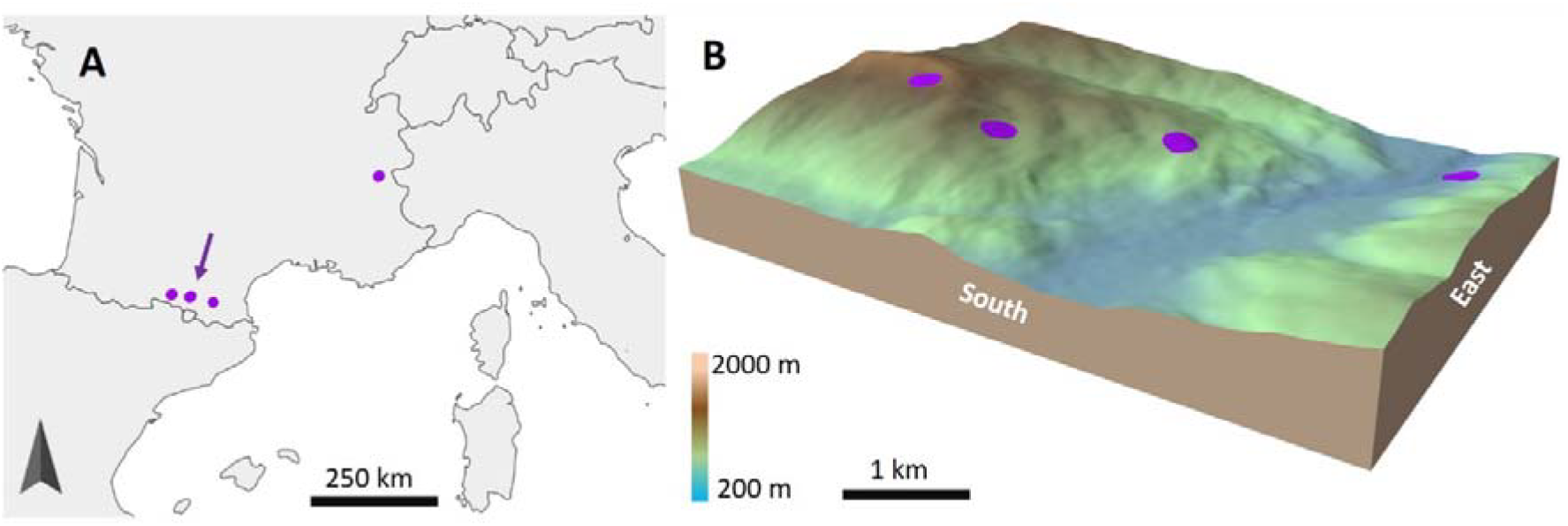
Sampling locations (A) and topographical zoom on the sampling location with arrow (B). See Tables 1 and S1 for topographical information on all populations.

**Table 1.**
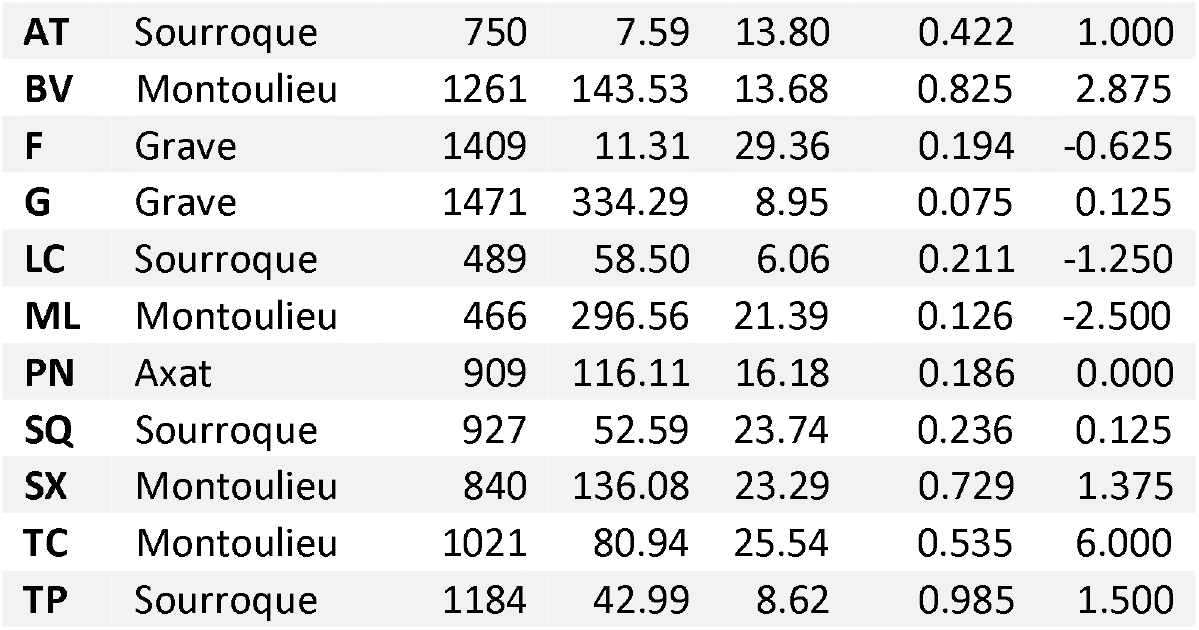
Sampling location features. NHeight (normalized height, 0-100%) reflects height relative to the nearest valley, and TPI (topographic position index, unitless) describes the curve in the landscape with TPI > 0 near ridges and TPI < 0 near valleys (see 2.4 Topography). Altitude is expressed as m a.s.l., and slope in percentages. Grave is the Alpine meta-population; all other meta-populations are Pyrenean.

### 2.2 Clonal propagation and common garden

After three weeks of cold treatment (°5C), seeds were germinated *in vitro* on sterile hormone free MS (Murashige & Skoog, 1962) medium. A total of five seedlings per mother plant were transplanted onto a meristem multiplication medium (MS supplemented with 0.5 mg/L BA (Benzyladenine)), which rendered ca. five shoot meristems per seedling. Each of these meristems was then transferred to new medium for a second multiplication cycle, aiming to obtain at least 15 clones per seedling (i.e. per genotype). These clones were allowed to grow and set roots on a rooting medium (MS supplemented with 1 mg/L IBA (Indole butyric acid)) until transplantation into individual pots with 0.45 L standard potting soil in February 2018. To maximize survival during acclimatization in greenhouse conditions, air humidity was kept at 90% for two weeks and then slowly decreased towards standard greenhouse conditions (ca. 70% humidity). Multiplication success, i.e. the average within genotype proportion of replicates surviving both the *in vitro* and acclimatization procedure, varied considerably among populations (from 31.6% in BV to 82.0% in G) (Table S2). We found a relatively strong correlation between multiplication success and vegetative propagation in the greenhouse (Pearson r = 0.59, see Fig S1), indicating a natural predisposition for clonal growth. No or weak correlations were found between multiplication success and other traits (Pearson r < 0.20, Fig S1), suggesting that the *in vitro* procedure did not considerably impact our results. A total of 1416 plants survived the acclimatization and were monitored throughout the experiment (population mean of 142, genotype mean of 12).

Pots were fully randomized across the greenhouse and across soil moisture treatments to minimize effects of micro-environmental variation on relations between drought treatments and phenotypic responses. Three soil moisture treatments (dry, moist, wet), which are assumed to capture at least part of the natural variation in soil moisture, were applied using a tube system (with 42 pots per tube) that allowed controlled watering (Fig S2). For a duration of two minutes, tubes were watered once or twice a week, depending on ambient temperatures (dry treatment), every (two) day(s) (moist treatment), and every four hours (wet treatment). Soil moisture was measured at two time points to quantify soil moisture differences between treatments, through measuring pot weights and through TDR (time-domain reflectometry) measurements. This indicated clear soil moisture differences among treatments (Fig S3). Starting in April 2018, the soil moisture treatments were applied during two cycles of six weeks, with a recuperation period of two weeks in between, in which all plants received equal amounts of water (cfr. moist treatment). During a third cycle of soil moisture treatments, far-red lamps (which mimic the light conditions typically experienced during flowering of *F. vesca* under tree cover) were installed to initiate flowering (Rantanen *et al.* 2014). We applied this treatment because, except for one population (“ML”), plants did not flower under regular greenhouse conditions.

### 2.3 Phenotypic traits and their plasticity

A total of five phenotypic traits were measured for all plants, namely dry above-ground biomass, specific leaf area for leaves without petiole (SLA for one leaf per plant, Pérez-Harguindeguy *et al.* 2013), the total number of runners produced during the second soil moisture treatment (counted and cut each week), stomatal density, and relative flowering rate (RFR) (Table S3). Stomata were counted on nail polish leaf prints using a KEYENCE light microscope at 1000 x magnification. For three replicate counts per leaf print, median stomatal counts instead of averages were used to minimize the impact of counting errors. **RFR** was calculated as (200 – t_ff_)/200, with t_ff_ = time of first flowering, expressed as days after initiation of the far-red treatment. Plants that did not flower after 100 days of monitoring were given a 200 score to obtain an RFR of zero. Plants originating from the “ML” population flowered prior to far-red treatment and were therefore excluded from the RFR analyses. To remove redundant variation among the different traits, a principal component analysis (PCA) was performed on the vegetative traits (above-ground biomass, SLA, number of runners and stomatal density). This resulted in three principal components (PCs) retaining 91.1% of the original variation, with PC1 (hereafter **Growth**) mainly representing above-ground biomass and number of runners (positively correlated), PC2 (hereafter **Stomata**) correlating (positively) with stomatal densities, and PC3 (hereafter **SLA**) correlating (negatively) with SLA (Table S3). PC3 was inverted so that it increases with increasing SLA, thereby facilitating interpretation.

The relative distance plasticity index (RDPI), which quantifies phenotypic distances among individuals exposed to different environments, has been proposed as the most appropriate plasticity index among 17 different plasticity metrics (Valladares et al. 2006). We determined RDPI for each trait (PC) and for each genotype as the absolute trait difference between all pairs of clonal replicates grown under the three distinct drought treatments divided by the phenotypic sum of the respective pairs (Valladares *et al.* 2006). RDPI ranges from 0 (no plasticity) to 1 (maximal plasticity), and was calculated using the R package *Plasticity (*Ameztegui 2017). This rendered the additional trait variables **Growth plasticity, Stomatal plasticity, SLA plasticity and RFR plasticity (Tables S4, S5)**.

### 2.4 Topography

A total of five topographical variables were extracted for each population using QGIS 2.18, namely aspect, slope (0-100%), elevation (m a.s.l.), normalized height (altitude relative to the nearest valley, 0-100%) and topographic position index (TPI). Aspect was decomposed into two independent variables describing orientation relative to the South direction (aspect_South) and orientation relative to the East direction (aspect_East). TPI describes the curve in the landscape, with high TPI (>0) near ridges (water efflux) and low TPI (<0) near valleys (water influx). Where no curve is present in the landscape (flat terrain and constant slopes), TPI reaches zero. The elevation map used for topographical extractions was at 30-meter resolution (gdex.cr.usgs.gov) and re-projected to a local coordinate reference system (Lambert Sud France). To obtain orthogonal topographical variables, a PCA was performed on all original topographical variables, and resulted in three PCs representing 83.6% of the variation across the original topographical variables (Table S1). The first PC (hereafter **Relative height**) was mainly shaped by TPI, normalized height and aspect_East, with low values representing populations in valleys with eastern aspect. The second PC (hereafter **Altitude**) was positively correlated with elevation and aspect_South, and the third PC (hereafter **Slope**) was positively correlated with slope. Note that “Altitude” reflects maximum daily temperature ranges as the south-oriented, high altitude populations receive most sun and therefore heat up considerably over the day while temperatures are low during the night as compared to low altitude populations that are north-oriented.

### 2.5 Modelling of traits and trait plasticity

**Growth, Stomata** and **SLA** were modeled using linear mixed models as implemented in the R package *lme4*. We accounted for genetic differentiation through the random intercept factor “Genotype”:

Trait ~ Relative height + Altitude + Slope + Treatment + Relative height × Treatment + Altitude × Treatment + Slope × Treatment + Meta-population + 1| Genotype

**Growth RDPI, Stomatal RPDI** and **SLA** RDPI across soil humidity treatments, are confined between 0 and 1 and were therefore modelled as beta distributions with logit link. Using the R package *glmmTMB (*Brooks *et al.* 2017), a generalized mixed model with a beta response distribution was built as follows:

Trait plasticity ~ Relative height + Altitude + Slope + Meta-population + 1| Genotype

Where required, response variables were transformed to improved model fits (see Table S6 for model residual plots). Pearson residuals, which are preferred when response variables follow a beta distribution (Espinheira *et al.* 2008), were used for evaluation of the generalized models (Table S7). Marginal (fixed model) and conditional (full model) R² values were computed using the R package *sjstats (*Lüdecke *2019)*.

To improve the fit of **RFR** (excess of zeros) and **RFR plasticity** (excess of ones) models, genotype averages were computed and modeled using beta regression as implemented in R package *betareg*. This package also provides a pseudo-R² value as an overall effect size for beta regressions. To correct for increased Type I error due to multiple hypothesis testing, p-values were adjusted using the false discovery rate approach implemented in the R package “*qvalue*” (Storey *et al.* 2019).

### 2.6 Heritability

For each treatment, heritability in the broad sense (H^2^) was calculated as V_G_/(V_G_ + V_E_/r), with V_G_ representing the variance between genotypes, V_E_ the residual variance (i.e. random environmental variation among clones), and r the number of replicates (clones) (Lynch & Walsh 1998). Variance components and H^2^ confidence intervals were obtained using a random effects model with the delta method implemented in the pin function of R package *sommer (*Covarrubias-Pazaran *2018)*. Trait H^2^ estimates at the population level were highly inaccurate (high SE), but were provided in Table S8 for completeness. We also calculated H^2^ at the population level for each of the plasticity traits, and plotted plasticity H^2^ against topographical PCs.

## 3 RESULTS

Drought significantly reduced growth, SLA and RFR compared to moist and wet treatments, while stomatal densities were significantly higher under drought stress than under moist and wet treatments (Figs. 2 and 3A, Table 2). The magnitude of these soil moisture treatment effects depended on the topographical features associated with the sampling locations of the populations, as demonstrated by significant interactions between treatment and at least one topographic variable (Table 2). Specifically, populations from low altitude and from high relative height and steep slopes experienced less growth reductions under drought stress than populations from high latitude, low relative height and weak slopes. In addition, populations from steep slopes retained high SLA under drought stress as opposed to populations from weak slopes (Fig. 2). The increase in stomatal densities with drought stress was most pronounced for populations from high altitude (Fig. 2). RFR did not vary significantly between populations from different topographical settings (Table 2).

**Fig. 2.**
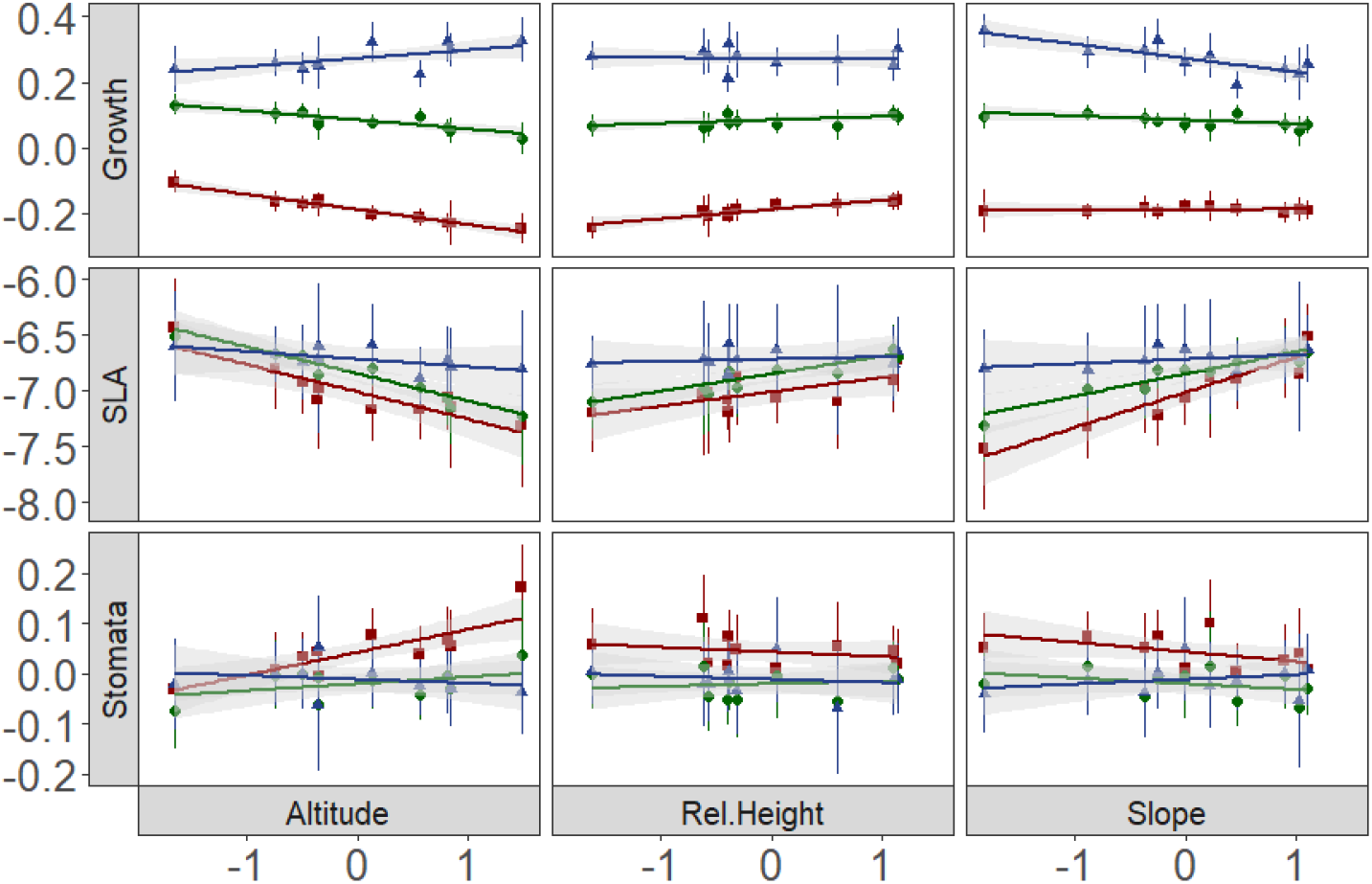
Relations between vegetative traits (principal components) and topographical variables (principal components) under three soil moisture treatments. Red squares, green dots and blue triangles represent dry, moist and wet soil moisture treatments, respectively. Plots present partial effects of the respective traits and their interaction with soil moisture treatment, extracted using the R package remef (Hohenstein 2013).

**Fig. 3.**
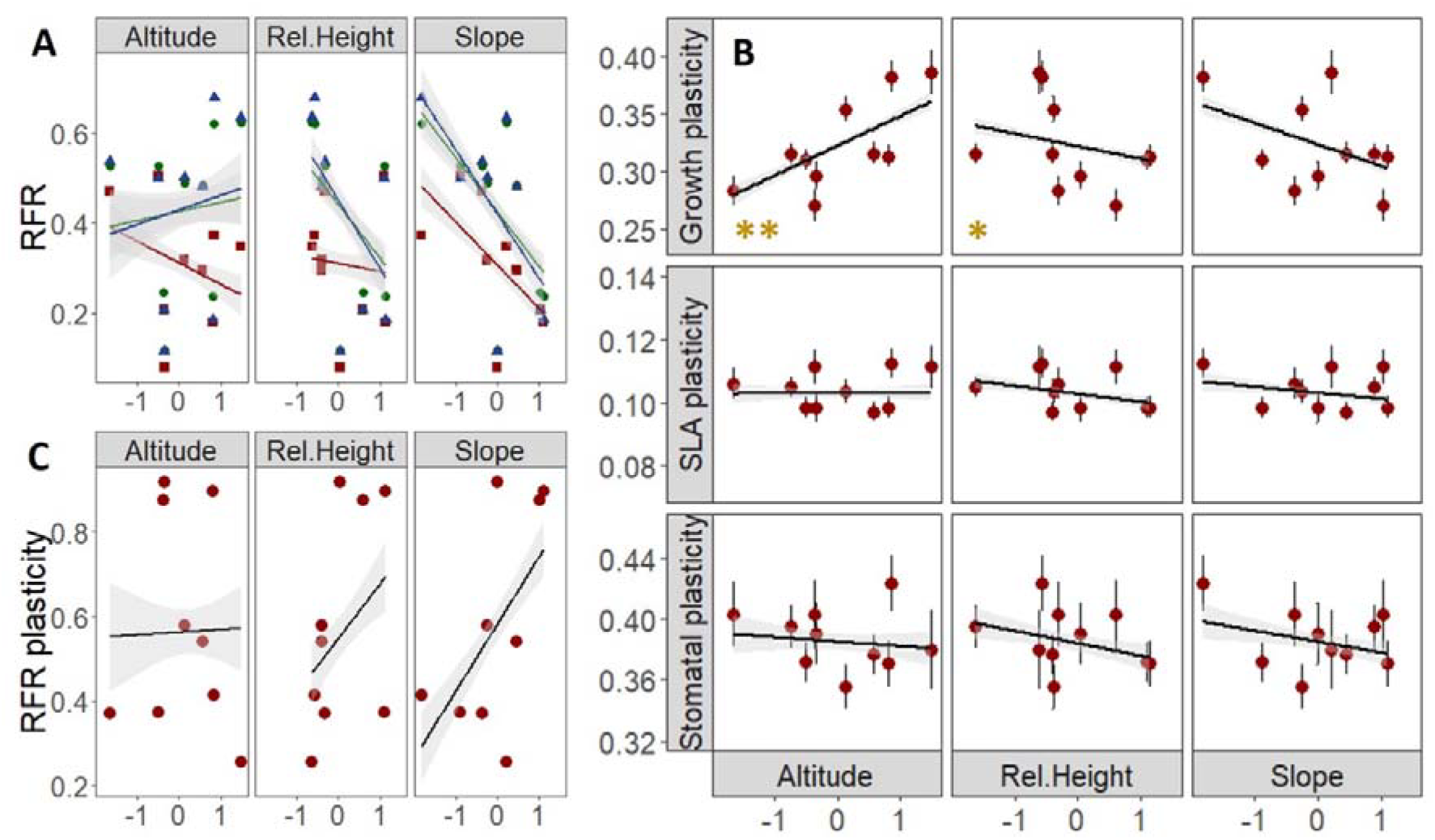
Relations between traits and topographical variables (principal components). (A) RFR against topography under three soil moisture levels, (B) vegetative trait plasticities against topography, and (C) RFR plasticity against topography. Red squares, green dots and blue triangles represent dry, moist and wet soil moisture treatments, respectively. Yellow asterisks represent the significance level for plasticity traits (* and ** are p-values <0.05 and <0.01, resp.)

**Table 2.** Significance of model terms, with *, ** and *** representing FDR corrected p-values < 0.05, < 0.01 and < 0.001, respectively. R^2^ represents R^2^_marginal_ for the mixed models, and pseudo-R^2^ for the beta regressions. See Tables S6 and S7 for detailed test statistics. NA = not applicable, NS = not significant (p>0.05).

| | Treatment | Rel.Height<sup>(xTreatment)</sup> | Altitude<sup>(xTreatment)</sup> | Slope<sup>(xTreatment)</sup> | $R^2$ |
| --- | --- | --- | --- | --- | --- |
| Growth | *** | *(*) | **(***) | NS(***) | 0.66 |
| Stomata | *** | NS<sup>(NS)</sup> | *(*) | NS<sup>(NS)</sup> | 0.06 |
| SLA | *** | NS<sup>(NS)</sup> | *(NS) | *(**) | 0.08 |
| RFR | ** | NS<sup>(NS)</sup> | NS<sup>(NS)</sup> | NS<sup>(NS)</sup> | 0.34 |
| Growth plasticity | NA | * | ** | NS | 0.15 |
| Stomatal plasticity | NA | NS | NS | NS | 0.08 |
| SLA plasticity | NA | NS | NS | NS | 0.01 |
| RFR plasticity | NA | NS | NS | NS | 0.36 |

Altitude patterns systematically and markedly mirrored relative height and slope patterns across traits and treatments (Figs. 2 and 3A), indicating that these topographical variables independently, but predictably affect plant responses to soil moisture treatments. More specifically, whereas growth under drought stress decreased with increasing latitude, it increased for populations situated near mountain summits (Fig. 2). Similarly, drought stress decreased SLA for populations originating from higher altitude, while it increased SLA for populations from relatively steep slopes.

Trait plasticity was higher for RFR (between 0.2 and 1.0) than for all vegetative traits (between 0.1 and 0.4), and could be explained by topography to a limited extent (Table 2, Fig. 3). Genotypes from low relative height and from high altitude in particular showed increased plasticity for growth (Fig. 3B). Plasticity for SLA, stomata and RFR, on the other hand, did not differ significantly between topographical settings (figs. 3B and 3C) (Table 2).

Trait H^2^ was highest for RFR and lowest for stomata (Table S8), indicating that RFR is under strong genetic control while stomatal variation is predominantly environmental (Fig. 4). Trait H^2^ was comparable among treatments (Fig. 4), suggesting that traits can evolve irrespective of soil moisture levels and thus in response to increased drought stress.

**Fig. 4.**
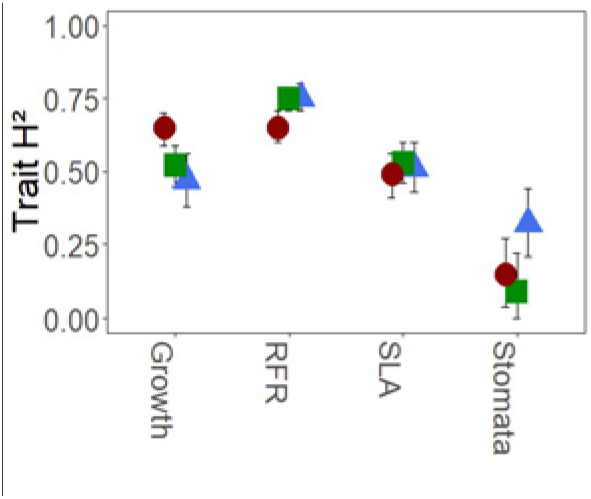
Trait heritability estimates across traits and soil moisture treatments. Red squares, green dots and blue triangles represent dry, moist and wet soil moisture treatments, respectively.

Similar to trait H^2^, plasticity H^2^ was on average highest for RFR (H^2^=0.62, SE=0.17), followed by growth (H^2^=0.48, SE=0.16), SLA (H^2^=0.46, SE=0.16) and stomata (H^2^=0.32, SE=0.17) (Table S8). For all traits, plasticity H^2^ tended to increase with increasing altitude (Fig. 5). Populations originating from higher altitudes may thus evolve increased trait plasticity when environmental conditions change. In addition, H^2^ of stomatal plasticity tended to decrease with increasing slope (Fig. 5).

**Fig. 5.**
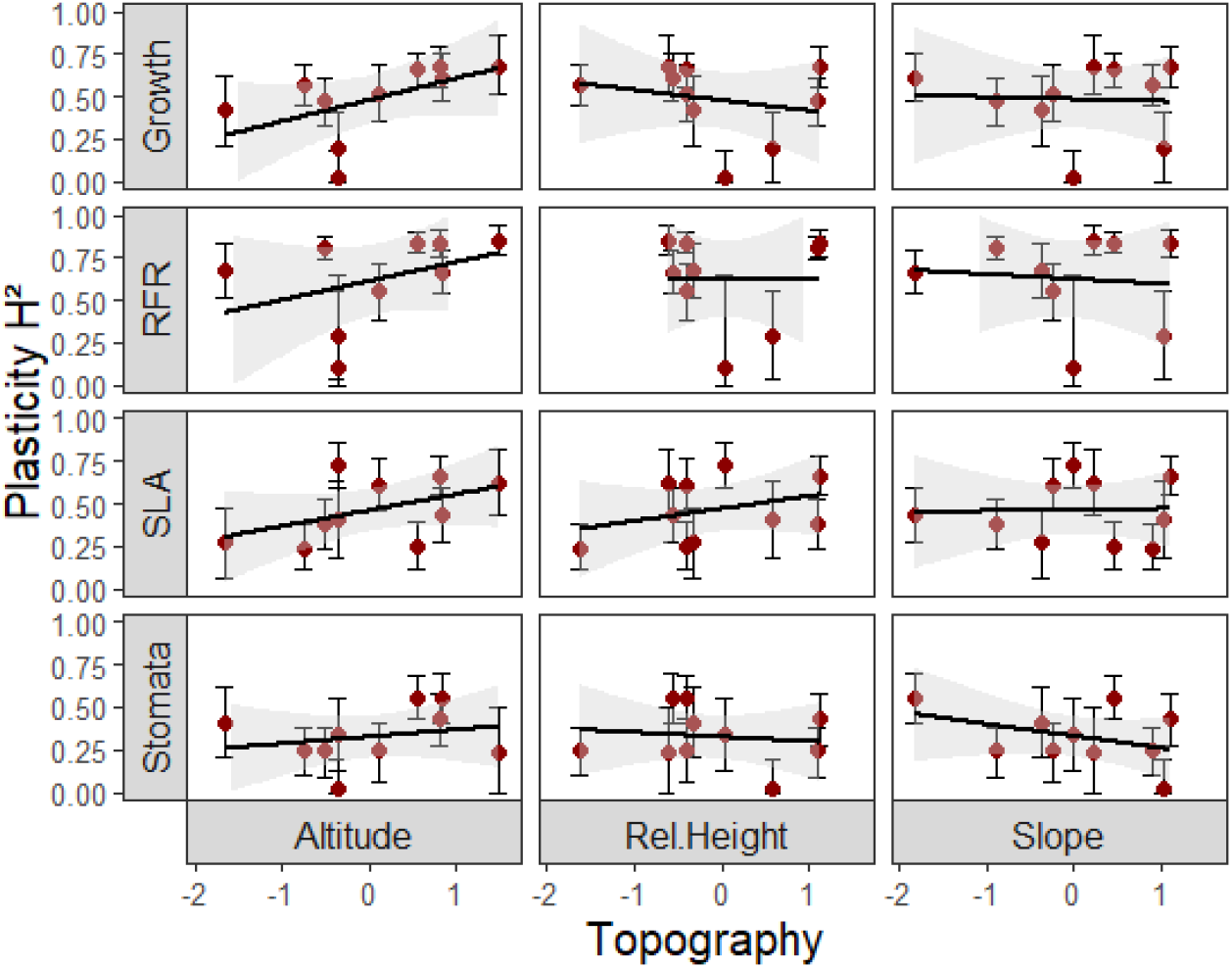
Plasticity heritability estimates across traits along topographical gradients.

## 4 DISCUSSION

Adaptive trait variation could mitigate the impact of changing soil moisture levels on population dynamics and persistence, yet little is known about the potential of mountainous populations to withstand increased levels of drought stress. The temporal soil moisture heterogeneity presumably associated to various topographical settings could boost adaptive evolution towards (i) phenotypes that increase drought resistance, and (ii) phenotypic plasticity for traits underlying drought resistance. We used genetically identical replicates (clones) of *Fragaria vesca*, originating from distinct topographical settings, to study whether topography can drive adaptive trait variation that allows population persistence under different levels of drought stress. Our results indicate that mountainous populations have a high potential to cope with drought stress, as they can evolve both through trait adaptation and through trait plasticity adaptation. However, low-altitudinal populations, which are expected to be most vulnerable to climate warming, have the lowest heritability and may thus only withstand modest increases in drought stress.

### 4.1 Genetic adaptive trait variation in mountainous populations

Plants adapted to dry environments should be able to maintain growth under increased levels of drought to a larger extent than plants originating from moist environments. We correspondingly found less growth reduction under drought stress for plants originating from higher relative height (~ water efflux) and low absolute altitude (~ high minimal temperatures) than for plants originating from cool and moist locations (Fig. 2). To maintain growth under drought stress, plants adapted to dry environments should retain photosynthetic rates at relatively high levels and thus minimize reductions in SLA (e.g. Wellstein et al. 2017). Accordingly, the plants originating from dry environments (steep slopes and high relative height) had significantly higher SLA under increased drought than plants originating from moist environments (weak slopes and valleys). In addition, plant populations that evolved to thrive in dry conditions have frequently been found to escape summer drought through advanced flowering time (Franks 2011; Nguyen et al. 2016). Here, although flowering tended to advance with increasing relative height and slope (~ water efflux), these trends were deemed non-significant (Table 2, Fig. 3A).

Stomatal densities increased significantly under drought stress (Fig. 2). An increase in stomatal density can result in an improved water use efficiency through indirect effects of stomatal size, as size and density usually are inversely correlated (Dittberner *et al.* 2018). Small stomata not only have lower transpiration rates, they can also open and close at higher velocity than large stomata in response to local soil moisture changes (Drake *et al.* 2013; Raven 2014). Plants adapted to dry environments may strongly benefit from high densities of small and responsive stomata, allowing quick re-initiation of photosynthesis when soil moisture levels temporarily increase. Here, plants from high absolute altitude in particular increased their stomatal density under drought (Fig. 2). High absolute altitude may correspond to decreased soil moisture levels where temperature inversions are frequent and/or where slopes are south-oriented. Our high-altitude populations were always south-oriented (Table S1), which may have driven evolution toward small stomata at high densities to deal with increased summer transpiration rates. Together, we could verify our hypothesis that genotypes originating from low altitude (~ high temperatures), high relative height and TPI (~ water efflux) and steep, southerly oriented slopes (~ water efflux and high evapotranspiration) have evolved traits that allow growth under drought stress. However, collinearity between altitude and aspect complicates disentangling the evolutionary roles of these two topographical in governing drought tolerance.

Although significant relations were observed between topography and phenotypic traits, much of the variation in SLA and stomatal density remained unexplained (Table 2, Table S6). Microclimatic variation not accounted for by the selected topographical variables (e.g. soil texture and shade) may have driven drought resistance evolution in addition to the selected topographical variables and thus explain some of the residual phenotypic variation. In addition, genetic drift likely contributes to the unexplained variation in phenotypic traits, as neutral processes are expected to shape at least part of the phenotypic variation observed in common gardens (McKay & Latta 2002; De Kort *et al.* 2013). We did not perform genetic marker analysis to exclude this neutral component (F_ST_, Wright 1943), but instead assumed topographical variables as likely drivers of drought-related trait variation to be plausible cues for adaptive evolution.

### 4.2 Adaptive variation in trait plasticity of mountainous populations

The use of genotypic replicates is key to studying adaptive divergence of phenotypic plasticity, which is predicted to drive evolutionary and demographic trajectories of natural populations. This study is among the first to quantify drought-related phenotypic plasticity as a trait that can vary between genotypes within populations and thus evolve over contemporary time scales. Increased plasticity has been suggested to provide a fitness advantage where environments are heterogeneous but predictable (Reed *et al.* 2010; Chevin & Lande 2011; Chevin & Hoffmann 2017; Bonamour *et al.* 2019). We found that within-genotype plastic responses to drought were common, but that only growth plasticity followed a pattern coinciding with topographical variation (Fig. 3B). Specifically, genotypes from low absolute altitude and from high relative height (near summits) were less plastic for growth. We suspect that increasing environmental heterogeneity with increasing absolute altitude, where south-facing populations experience fluctuating temperatures and soil moisture levels, may be an important driver for increased growth plasticity. In addition, environmental predictability may decrease away from valleys, thereby hampering evolution of phenotypic plasticity. Together, population inhabiting high altitudinal valleys seem to benefit most from high levels of growth plasticity.

The capacity of a genotype to adjust its gene expression in response to a change in environmental conditions is frequently referred to as epigenetic variation (Bossdorf *et al.* 2008; Thorson *et al.* 2017). Here, we thus provide indirect evidence that populations can evolve their epigenetic machinery towards increased epigenetic flexibility in environments with heterogeneous soil moisture levels. Although habitat differences have been shown to result in epigenetic divergence between populations (e.g. Lele et al. 2018, Schmid et al. 2018), empirical evidence for evolution toward higher within-population epigenetic variation in heterogeneous environments is still lacking.

### 4.3 Drought-related phenotypic plasticity has a substantial heritable component

A large proportion of the phenotypic variation could be attributed to genetic rather than environmental variation, as indicated by the relatively high heritability estimates. This genetic variation can serve as a basis for natural selection to act upon, and thus reflects the potential of populations to evolve under changing environmental conditions (Mousseau & Roff 1987; Visscher *et al.* 2008). A spatially varying balance between gene flow and natural selection pressures likely lies at the basis of this accumulation of adaptive genetic variation. First, steep fine-scale environmental variation can cause strong and spatially variable selection pressures that, in combination with gene flow-facilitated spread of adaptive genetic variants, maintain adaptive genetic variation (Hämälä *et al.* 2018; Schmidt & Garroway 2018). Second, opposing selection pressures from several environmental factors affecting the same phenotypic traits (e.g. growth can be affected by soil moisture levels, other abiotic factors as well as the local biotic context) promote the maintenance adaptive genetic variation (Kruuk *et al.* 2002, 2008; Alberto *et al.* 2011).

With heritability estimates fluctuating around 0.5 (Figs. 4 and 5), phenotypic traits were found to be comparably heritable as their plasticity indices, indicating that traits as well as phenotypic plasticity are under genetic control and can evolve in response to changes in soil moisture levels. Growth, flowering rate, and their corresponding plasticity indices showed particularly high heritability (Figs. 4 and 5), suggesting that these fitness traits have high potential to keep pace with local environmental changes (e.g. in pollinator phenology, drought levels and temperature). On the other hand, low-altitudinal populations harboured a considerably lower heritability than populations at high absolute altitude (Fig. 5). The higher environmental heterogeneity at high altitude may have facilitated the accumulation of heritable phenotypic variation, while the lack hereof at low altitude increases the vulnerability of low altitudinal populations to environmental change. We conclude that populations that evolved to thrive in heterogeneous conditions likely are pre-adapted to climate change. Such populations represent invaluable sources of quantitative genetic variation that could support conservation where climate projections are inconclusive.

## ACKNOWLEDGMENTS

This study was supported by the Research Foundation of Flanders (FWO). Multiplication of *F. Vesca* accessions was coordinated by Kevin Longin and Dr. Bart Panis. Transplantation of *F. vesca* seedlings from *in vitro* cultures to greenhouse settings took place with the help of Kasper Van Acker, Sophie Degrande, Eva Hulsman, Arne Sinnesael, Pablo Deschepper and Lore Geeraert. Kasper Van Acker, Jan Gerits and Gerrit Peeters also helped maintaining the greenhouse experiment. We thank Poi Verwilt for his support with the experimental set-up and greenhouse preparations. Christophe Coeck calibrated the TDR meter prior to drought measurements.

HDK samples all Pyrenean populations, generated the project, collected and analyzed the data, and wrote the first version of the manuscript. All co-authors provided comments on the manuscript. In addition, BP developed the *in vitro* protocol, KH provided crucial help in the greenhouse, RD collected seeds in the Alpine locations and SJ supported the microscopic processing of the leaf prints.

## DATA ARCHIVING

All data required for the data analysis are available in the supporting files.

## REFERENCES

Alberto, F., Bouffier, L., Louvet, J.-M., Lamy, J.-B., Delzon, S. & Kremer, a. (2011). Adaptive responses for seed and leaf phenology in natural populations of sessile oak along an altitudinal gradient. J. Evol. Biol., 24, 1442–54.

Alexander, J.M., Chalmandrier, L., Lenoir, J., Burgess, T.I., Essl, F., Haider, S., et al. (2018). Lags in the response of mountain plant communities to climate change. Glob. Chang. Biol., 24, 563–579.

Ameztegui, A. (2017) Plasticity: An R package to determine several plasticity indices.

Anderson, J.T., Willis, J.H. & Mitchell-Olds, T. (2011). Evolutionary genetics of plant adaptation. Trends Genet., 27, 258–66.

Arnold, P.A., Kruuk, L.E.B. & Nicotra, A.B. (2019). How to analyse plant phenotypic plasticity in response to a changing climate. New Phytol., 222, 1235–1241.

Bansal, S., Harrington, C.A., Gould, P.J. & St.Clair, J.B. (2015). Climate-related genetic variation in drought-resistance of Douglas-fir (Pseudotsuga menziesii). Glob. Chang. Biol., 21, 947–958.

Bertrand, R., Riofrío-Dillon, G., Lenoir, J., Drapier, J., de Ruffray, P., Gégout, J.-C., et al. (2016). Ecological constraints increase the climatic debt in forests. Nat. Commun., 7, 12643.

Bonamour, S., Chevin, L.-M., Charmantier, A. & Teplitsky, C. (2019). Phenotypic plasticity in response to climate change: the importance of cue variation. Philos. Trans. R. Soc. B Biol. Sci., 374, 20180178.

Bossdorf, O., Richards, C.L. & Pigliucci, M. (2008). Epigenetics for ecologists. Ecol. Lett., 11, 106–15.

Brooks, M.E. et al. (2017). glmmTMB Balances Speed and Flexibility Among Packages for Zeroinflated Generalized Linear Mixed Modeling. The R Journal, 9(2), 378–400.

Chevin, L.-M. & Hoffmann, A.A. (2017). Evolution of phenotypic plasticity in extreme environments. Philos. Trans. R. Soc. B Biol. Sci., 372, 20160138.

Chevin, L.-M. & Lande, R. (2011). Adaptation to marginal habitats by evolution of increased phenotypic plasticity. J. Evol. Biol., 24, 1462–76.

Cooper, H.F., Grady, K.C., Cowan, J.A., Best, R.J., Allan, G.J. & Whitham, T.G. (2019). Genotypic variation in phenological plasticity: Reciprocal common gardens reveal adaptive responses to warmer springs but not to fall frost. Glob. Chang. Biol., 25, 187–200.

Covarrubias-Pazaran, G. (2018). Software update: Moving the R package sommer to multivariate mixed models for genome-assisted prediction.

Dittberner, H., Korte, A., Mettler-Altmann, T., Weber, A.P.M., Monroe, G. & de Meaux, J. (2018). Natural variation in stomata size contributes to the local adaptation of water-use efficiency in Arabidopsis thaliana. Mol. Ecol., 27, 4052–4065.

Drake, P.L., Froend, R.H. & Franks, P.J. (2013). Smaller, faster stomata: scaling of stomatal size, rate of response, and stomatal conductance. J. Exp. Bot., 64, 495–505.

Espinheira, P.L., Ferrari, S.L.P. & Cribari-Neto, F. (2008). On beta regression residuals. J. Appl. Stat., 35, 407–419.

Günther, T., Lampei, C., Barilar, I. & Schmid, K.J. (2016). Genomic and phenotypic differentiation of Arabidopsis thaliana along altitudinal gradients in the North Italian Alps. Mol. Ecol., 25, 3574–3592.

Halbritter, A.H., Fior, S., Keller, I., Billeter, R., Edwards, P.J., Holderegger, R., et al. (2018). Trait differentiation and adaptation of plants along elevation gradients. J. Evol. Biol., 31, 784–800.

Hämälä, T., Mattila, T.M. & Savolainen, O. (2018). Local adaptation and ecological differentiation under selection, migration, and drift in Arabidopsis lyrata *. Evolution, 72, 1373–1386.

Hoffmann, A.A., Sgrò, C.M. & Kristensen, T.N. (2017). Revisiting Adaptive Potential, Population Size, and Conservation. Trends Ecol. Evol., 32, 506–517.

Hohenstein (2013). The remef function for R.

IPCC. (2014). and and. Clim. Chang. 2014 Synth. Report. Contrib. Work. Groups I, II III to Fifth Assess. Rep. Intergov. Panel Clim. Chang. Pachauri RK, Meyer LA, eds. Geneva, Switz. IPCC.

Kingsolver, J.G. & Buckley, L.B. (2017). Evolution of plasticity and adaptive responses to climate change along climate gradients. Proc. R. Soc. B Biol. Sci., 284, 20170386.

van Kleunen, M. & Fischer, M. (2005). Constraints on the evolution of adaptive phenotypic plasticity in plants. New Phytol., 166, 49–60.

Körner, C. (2007). The use of ‘altitude’ in ecological research. Trends Ecol. Evol., 22, 569–574.

De Kort, H., Vander Mijnsbrugge, K., Vandepitte, K., Mergeay, J., Ovaskainen, O. & Honnay, O. (2016). Evolution, plasticity and evolving plasticity of phenology in the tree species Alnus glutinosa. J. Evol. Biol., 29, 253–264.

De Kort, H., Vandepitte, K. & Honnay, O. (2013). A meta-analysis of the effects of plant traits and geographical scale on the magnitude of adaptive differentiation as measured by the difference between QST and FST. Evol. Ecol., 27, 1081–1097.

Kruuk, L.E.B., Slate, J., Pemberton, J.M., Brotherstone, S., Guinness, F. & Clutton-Brock, T. (2002). Antler size in red deer: heritability and selection but no evolution. Evolution, 56, 1683–1695.

Kruuk, L.E.B., Slate, J. & Wilson, A.J. (2008). New Answers for Old Questions: The Evolutionary Quantitative Genetics of Wild Animal Populations. Annu. Rev. Ecol. Evol. Syst., 39, 525–548.

Lele, L., Ning, D., Cuiping, P., Xiao, G. & Weihua, G. (2018). Genetic and epigenetic variations associated with adaptation to heterogeneous habitat conditions in a deciduous shrub. Ecol. Evol., 8, 2594–2606.

Lüdecke, D. (2019). sjstats: Statistical Functions for Regression Models (Version 0.17.4).

Lynch, M. & Walsh, B. (1998). Genetics and analysis of quantitative traits. Sunderland, MA: Sinauer.

McKay, J.K. & Latta, R.G. (2002). Adaptive population divergence: markers, QTL and traits. Trends Ecol. Evol., 17, 285–291.

Mousseau, T.A. & Roff, D.A. (1987). Natural selection and the heritability of fitness components. Heredity, 59, 181–197.

Murashige, T. & Skoog, F. (1962). “A Revised Medium for Rapid Growth and Bio Assays with Tobacco Tissue Cultures”. Physiologia Plantarum. 15 (3): 473–497.

O’Brien, E.K., Higgie, M., Reynolds, A., Hoffmann, A.A. & Bridle, J.R. (2017). Testing for local adaptation and evolutionary potential along altitudinal gradients in rainforest Drosophila[]: beyond laboratory estimates. Glob. Chang. Biol., 23, 1847–1860.

Palacio-López, K., Beckage, B., Scheiner, S. & Molofsky, J. (2015). The ubiquity of phenotypic plasticity in plants: a synthesis. Ecol. Evol., 5, 3389–3400.

Pérez-Harguindeguy et al. (2013). New handbook for stand-ardised measurement of plant functional traits worldwide. Aus-tralian Journal of Botany, 61, 167–234.

Pfennigwerth, A.A., Bailey, J.K. & Schweitzer, J.A. (2017). Trait variation along elevation gradients in a dominant woody shrub is population-specific and driven by plasticity. AoB Plants, 9.

Rantanen, M., Kurokura, T., Mouhu, K., Pinho, P., Tetri, E., Halonen, L., et al. (2014). Light quality regulates flowering in FvFT1/FvTFL1 dependent manner in the woodland strawberry Fragaria vesca. Front. Plant Sci., 5, 271.

Raven, J.A. (2014). Speedy small stomata? J. Exp. Bot., 65, 1415–1424.

Reed, T.E., Waples, R.S., Schindler, D.E., Hard, J.J. & Kinnison, M.T. (2010). Phenotypic plasticity and population viability: the importance of environmental predictability. Proc. Biol. Sci., 277, 3391–400.

Scheepens, J.F., Deng, Y., Bossdorf, O. (2018) Phenotypic plasticity in response to temperature fluctuations is genetically variable, and relates to climatic variability of origin, in Arabidopsis thaliana. Annals of Botany Plants, 10(4), doi.org/10.1093/aobpla/ply043.

Schmid, M.W., Heichinger, C., Coman Schmid, D., Guthörl, D., Gagliardini, V., Bruggmann, R., et al. (2018). Contribution of epigenetic variation to adaptation in Arabidopsis. Nat. Commun., 9, 4446.

Schmidt, C. & Garroway, C.J. (2018). Digest: Local adaptation at close quarters*. Evolution, 72, 1531–1532.

Storey, J., Bass, A., Dabney, A. & Robinson, D. (2019). qvalue: Q-value estimation for false discovery rate control. R package version 2.14.1.

Sultan, S.E. (2001) Pphenotypic plasticity for fitness components in Polygonum species of contrasting ecological breadth. Ecology, 82(2), 328–343.

Thorson, J.L.M., Smithson, M., Beck, D., Sadler-Riggleman, I., Nilsson, E., Dybdahl, M., et al. (2017). Epigenetics and adaptive phenotypic variation between habitats in an asexual snail. Sci. Rep., 7, 14139.

Urban, M.C., Malcolm, J.R., Liu, C., Neilson, R.P., Hansen, L., Hannah, L., et al. (2015). Climate change. Accelerating extinction risk from climate change. Science, 348, 571–3.

Valladares, F., Sanchez-Gomez, D. & Zavala, M.A. (2006). Quantitative estimation of phenotypic plasticity: bridging the gap between the evolutionary concept and its ecological applications. J. Ecol., 94, 1103–1116.

Vázquez, D.P., Gianoli, E., Morris, W.F. & Bozinovic, F. (2017). Ecological and evolutionary impacts of changing climatic variability. Biol. Rev., 92, 22–42.

Visscher, P.M., Hill, W.G. & Wray, N.R. (2008). Heritability in the genomics era — concepts and misconceptions. Nat. Rev. Genet., 9, 255–266.

Waterhouse, M.D., Erb, L.P., Beever, E.A. & Russello, M.A. (2018). Adaptive population divergence and directional gene flow across steep elevational gradients in a climate-sensitive mammal. Mol. Ecol., 27, 2512–2528.

Wellstein, C., Poschlod, P., Gohlke, A., Chelli, S., Campetella, G., Rosbakh, S., et al. (2017). Effects of extreme drought on specific leaf area of grassland species: A meta-analysis of experimental studies in temperate and sub-Mediterranean systems. Glob. Chang. Biol., 23, 2473–2481.

Wilczek, A.M., Burghardt, L.T., Cobb, A.R., Cooper, M.D., Welch, S.M. & Schmitt, J. (2010). Genetic and physiological bases for phenological responses to current and predicted climates. Philos. Trans. R. Soc. B Biol. Sci., 365, 3129–3147.

Wright, S. (1943). Isolation by distance. Genetics, 28, 139–156.

Zhang, J.-J., Fischer, M., Colot, V., Bossdorf, O. (2012) Epigenetic variation creates potential for evolution of plant phenotypic plasticity. New Phytologist, 197(1), 314–322.

