## Supplementary material for "Pre-adaptation to climate change through topography-driven evolution of traits and their plasticity": Fig S

**Supplemental figures**

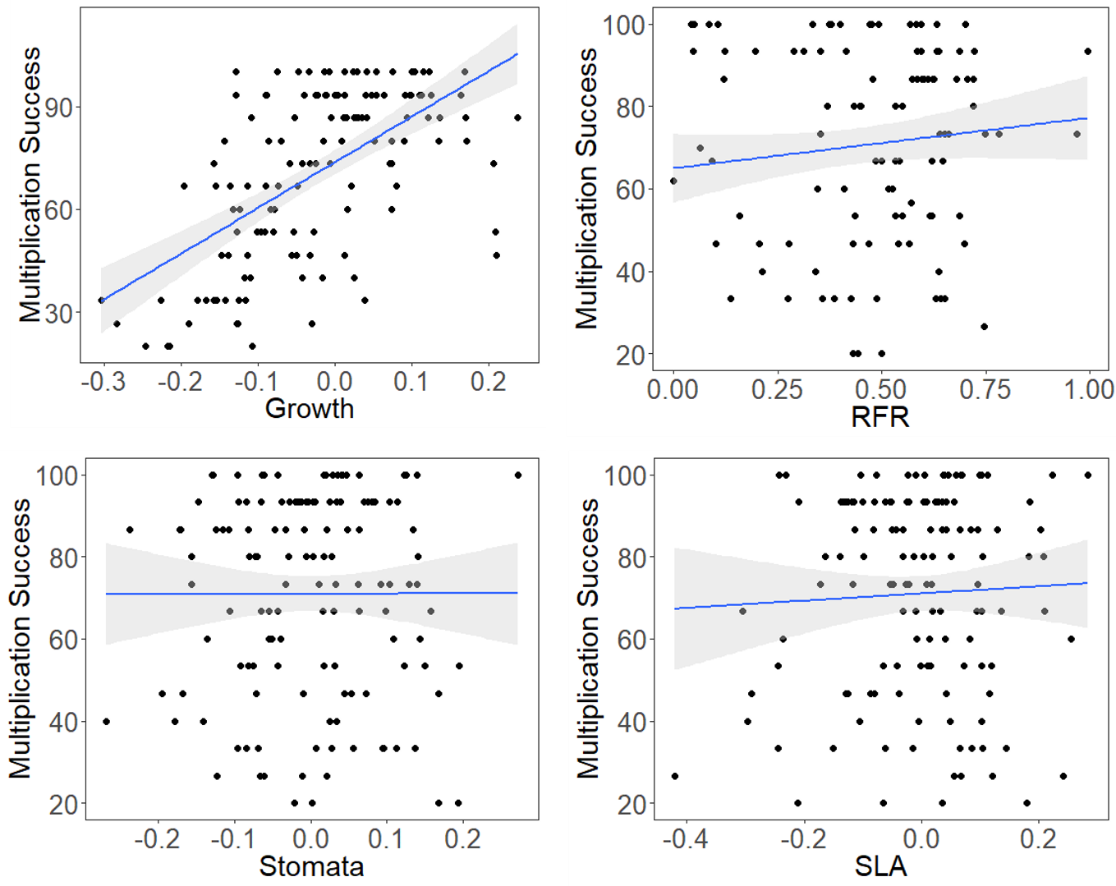

**Fig. S1.** Correlations between multiplication success (proportion survival after in vitro multiplication and transplantation to greenhouse).

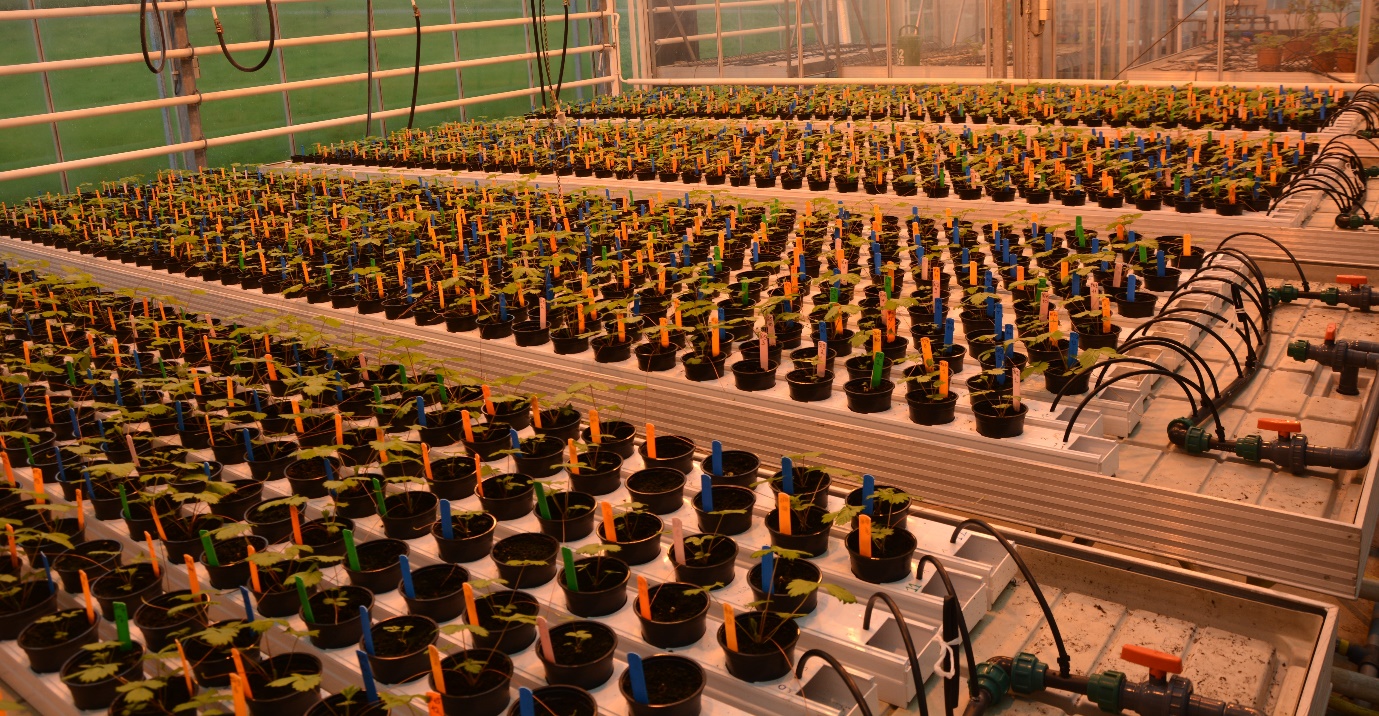

**Fig. S2.** Greenhouse set-up at the start of the experiment. Pots were fully randomized across four greenhouse tables (label colors correspond to metapopulations). Water administration occurred automatically through small black tubes, which were bent off manually during drought stress periods for a subset of the plant rows. A second set of water tubes was added for the subset of plant rows that received an excess of water (not yet installed here).

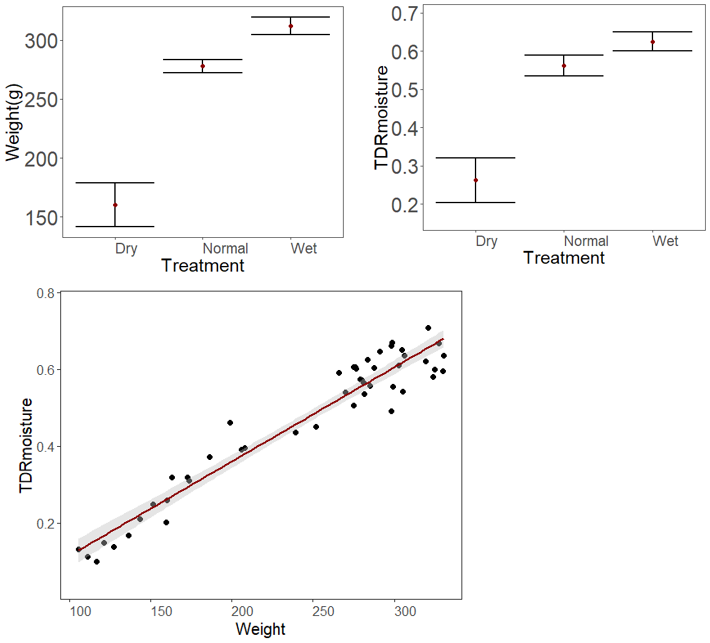

**Fig. S3.** Soil moisture differences among treatments as indicated by pot weight and TDR measurements.
